# Bacteroidia and Clostridia genomes collectively encode for a progressive cascade of marine polysaccharide degradation along the hindgut of the herbivorous fish *Kyphosus sydneyanus*

**DOI:** 10.1101/2023.12.23.570891

**Authors:** Cesar T. Facimoto, Kendall D. Clements, W. Lindsey White, Kim M. Handley

## Abstract

The gut microbiota of the marine herbivorous fish *Kyphosus sydneyanus* are thought to play an important role in host nutrition by supplying short-chain fatty acids (SCFA) through fermentation of dietary macroalgae. Here, we assembled 645 metagenome-assembled genomes (MAGs) from wild fish to determine the capacity of different bacterial taxa to degrade seaweed carbohydrates along the gut. Most bacteria (99%) were unclassified at the species level, highlighting taxonomic novelty dominated by Bacteroidia and Clostridia within the gut community. The presence of genes encoding endo-acting CAZymes in both phyla suggest they have a role in initiating glycan depolymerization. Bacteroidia also contributed the most to CAZyme-related gene expression in the distal hindgut, and encoded the highest densities of CAZymes within the community. In particular, the enrichment of CAZyme gene clusters (CGCs) within the Bacteroidia genus *Alistipes* (n = 73 versus just 59 distributed across all other taxa) points to an enhanced capacity for macroalgal polysaccharide utilization (e.g., alginate, laminarin and sulfated polysaccharides). Pairwise correlations of MAG relative abundances and encoded CAZyme compositions provide evidence of potential inter-species collaborations, whereby co-abundant MAGs exhibited complementary degradative capacities for specific substrates. Results indicated flexibility across these co-abundant groups in their capacity to source carbon (e.g., glucose or galactose-rich glycans), which possibly facilitates coexistence via niche partitioning. Our results indicate the potential for collaborative microbial carbohydrate metabolism in the gut of *K. sydneyanus* by Bacteroidia and Clostridia, and suggest that members of the genus *Alistipes* are a metabolically and taxonomically diverse group of specialized macroalgae biomass degraders.

## INTRODUCTION

Some marine herbivorous fishes rely on symbiotic relationships with microbial gut communities for nutrition and health [1, 2]. Significant levels of fermentation products are found within the gut of herbivorous taxa such as kyphosid chubs and unicornfishes of the genus *Naso*, and turnover rates of the resulting short-chain fatty acids (SCFA) are comparable to those in herbivorous mammals and reptiles [1]. These factors emphasize the importance of microbial fermentation in supporting the energetic metabolism of the fish host. In particular, the family Kyphosidae exhibits the highest levels of SCFA measured to date [3, 4]. Of the Kyphosidae, *Kyphosus sydneyanus* (Silver Drummer) displays exceptional levels of SCFA [1], highlighting a significant role for microbial gut communities in this species.

The microbial fermentation in the *K. sydneyanus* gut is fuelled by a mixture of carbohydrates derived from brown and red algae that are recalcitrant to host enzymes [2, 5, 6]. Investigation of the stomach contents of *K. sydneyanus* has revealed a prevalence of the brown algae species *Carpophylum maschalocarpum* (constituting 10-50% of stomach content) and *Ecklonia radiata* (15-50% of stomach content) [2]. Additionally, *Gigartina macrocarpa* (5-30% of stomach content) and *Caulacanthus ustulatus* (up to 15% of gut content) represent common red algae species present [2]. Therefore, the gastrointestinal tract of *K. sydneyanus* is enriched with a diverse array of carbohydrates such as alginate, laminarin, mannitol, and fucose-containing sulfated polysaccharides (FCSP) present in brown algae species. In addition, the gut contains carrageenan, agarose and floridean starch from dietary red algae [5, 7–9]. These substrates likely constitute the bulk of fermentation substrates supplied to gut microbial communities and govern metabolism along the fish gut.

The *K. sydneyanus* gut microbial composition has been explored recently, illustrating the predominance of Bacteroidota and Bacillota phyla across 60 individuals [10]. However, the relative abundances of these phyla differ longitudinally along the fish gut from mid-to distal sections [10, 11]. Certain families within the Bacillota, such as *Lachnospiraceae*, *Erysipelatoclostridiaceae*, *Oscillospiraceae* and *Acholeplasmataceae*, are significantly more abundant in the midgut [10, 12]. In contrast, the phylum Bacteroidota exhibits a gradual increase in relative abundance from mid-to distal gut, and is primarily represented by the family *Rikenellaceae* [10–12]. These observed variations in community composition across different gut sections, along with differing levels of SCFA, indicate regional variation in gut metabolism [12]. Despite longitudinal shifts in microbial composition, metagenomic investigation of the *K. sydneyanus* hindgut using un-binned contigs indicated the conservation of metabolic pathways along the gut [11], including redundancy in the metabolic capacity for algal degradation. Further metagenomic analysis of other *Kyphosus* species (e.g., *K. vaigiensis*, *K. hawaiiensis*, and *K. cinerascens*) revealed the enrichment of agarases, porphyranases, carrageenases, ulvanases and alginate-lyases primarily supplied by Bacteroidota [13]. How this capacity is distributed among the hindgut microbiota of *K. sydneyanus* is yet to be determined.

Here, we investigated the distribution of carbohydrate degradation capacity among the *K. sydneyanus* hindgut microbiota. We generated metagenome-assembled genomes (MAGs) and metatranscriptomes from different longitudinal sections of the lumen hindgut. Next, we determined the carbohydrate active enzyme (CAZyme) repertoire encoded by these genomes, and the capacity to break-down polysaccharides, particularly those comprising the *K. sydneyanus* diet of brown (e.g., *C. maschalocarpum* and *E. radiata*) and red algae (e.g., *G. macrocarpa*) [2]. This included the polysaccharides alginate, laminarin and FCSP that are important constituents of brown algae [7, 8, 14, 15], as well as carrageenan, the major cell wall polysaccharide in *Gigartina* species [9]. This diverse and complex substrate landscape, stemming from the range of highly polymerized and decorated polysaccharides (heteropolysaccharides) in the fish diet, requires a complex cascade of enzymatic processes, which in turn demands greater genetic diversity than is present in a single organism [16]. Accordingly, our findings show that multiple microbial consortia carry broadly analogous carbohydrate utilization capacities. Of these, Bacteroidia (particularly *Alistipes*) and Clostridia harbour endo-acting enzymes, indicating they are the main drivers of glycan degradation. These findings shed light on potential cooperative dynamics within the gut microbiota, and suggest a key role for Bacteroidia in initiating glycan degradation.

## MATERIALS AND METHODS

### Sample collection, metagenomic and metatranscriptomic sequencing

Four fish were collected in the summer of 2017 by underwater spear on snorkel from waters adjacent to the neighbouring Great Barrier and Little Barrier islands in north-eastern New Zealand. An additional six fish were collected via the same method in summer 2020 from Great Barrier Island. The initial four fish were used for generating metagenomes, and all analyses relate to these four fish unless otherwise specified. The six fish from 2020 were used for metagenomics and metatranscriptomics, and metagenomic results between the 2017 and 2020 are compared to illustrate the high reproducibility of results across individual fish and sampling years (Figures S1, S2). Fish collections from 2017 and 2020 were covered by approvals 001636 and 001949 from the University of Auckland Animal Ethics Committee. Gut sampling for metagenomic and metatranscriptomic sequencing was conducted as described previously [11]. To summarise, the gastrointestinal tract of each fish was separated immediately after capture and divided into sections [1]. The stomach was assigned as section I and the hindgut chamber as section V. The remaining intestine was divided into three sections of equal length and designated as II (immediately after the stomach), III and IV (immediately before the hindgut sphincter).

DNA from the gut lumen contents of sections IV and V was obtained from all ten fish caught in 2017 and 2020. Lumen contents from section III was also collected from the six 2020 fish. All eight samples from 2017 [11] and eighteen from 2020 were used for metagenomics. RNA was extracted from the lumen contents of the six 2020 fish based on detectable RNA of good quality (RIN values >5): all section V samples, three for section III, and four for section IV. Metagenome and metatranscriptome generation, MAG reconstruction, and transcript mapping to MAGs are described in Supplementary Information.

### Taxonomy and phylogenetic tree

Taxonomic classification of dereplicated MAGs (sharing <99% average nucleotide identity, Supplementary Information) was undertaken using the Genome Taxonomic Database Toolkit (GTDB-Tk) version 2.1.1 with the 214 database release [17]. A phylogenetic tree was produced with the protein sequence alignment file provided by GTDB-Tk and with IQ-TREE version 1.6.12 (parameters -m TEST -b 1000) [18] and reference genomes from GTDB.

### Gene prediction and annotation

Genes were predicted with PRODIGAL version 2.6.3 using meta mode [19]. Protein sequences were annotated using USEARCH version 9.02132 [20] against UniRef100 release 2018_09 [21], UniProt (SwissProt and TrEMBL) release 2018_09 [22], and KEGG release 86.1 [23] databases, considering best hits with at least 50% with reference coverage by length, 30% identity, and an e-value of 0.001. A protein domain search using HMMER v3.1b2 [24] was performed against Pfam version 32.0 [25] and TIGRFAM version 14.0 [26] (0.001 e-value cutoff).

CAZymes were annotated using dbCAN version 3.0.7 [27] using the protein mode and outputting CGCs. A CAZyme annotation was considered correct if at least two of the dbCAN output annotations (e.g., HMMER, eCAMI or DIAMOND) matched. The final CAZyme annotation prioritized HMMER annotations followed by eCAMI. The inference of CGCs substrate specificities were based on the presence of dedicated CAZy families and/or their colocalization with CAZy families containing complementary activities (Supplementary Information).

A BLAST-like DIAMOND search was also performed against the SulfAtlas version 2.3.1 database [28], and a subset of proteins from CAZy database version 3.0.7 [27] containing only proteins assigned to an EC number. The search was performed with DIAMOND version 2.0.15 [29] using blastp mode with a e-value cut off of 0.001. The annotation of ECs associated to degradation of brown and red algae polysaccharides are detailed in Supplementary Information. Sulfatases were only considered if the database target coverage was ≥50%, sequence identity ≥30%, and annotations contained the protein domain “PF00884” [30]. CAZymes associated with an EC were only considered if the database target coverage was ≥40% and identity ≥30%. Details about the curation of mannitol associated genes are in Supplementary Information.

### Statistical analysis and average nucleotide identity calculations

Statistical analyses were carried out with R version 4.1.1 [31]. A Wilcoxon rank-sum test was performed to determine CAZyme density across class using the ggpubr package [32]. Bray-Curtis dissimilarities were constructed using the vegan package [33] with MAG CAZyme densities. The package rstatix [34] was used to calculate statistical differences in relative abundance of genera across sections, and average predicted number of CGCs per genus, using the Kruskal-Wallis test. To identify clusters of co-abundant MAGs, pairwise Pearson correlation coefficients of MAG relative abundances were calculated using the rcorr function from the Hmisc package [35] and the heatmap was generated using corrplot [36]. Average Nucleotide Identities (ANI) between *A. putredinis*, *A. communis* and *Alistipes* MAGs from this study were calculated using the ANI calculator (http://enve-omics.ce.gatech.edu/ani/).

## RESULTS AND DISCUSSION

### Bacteroidia and Clostridia are major CAZyme-encoding class in *K. sydneyanus* hindgut

A total of 197 MAGs were recovered from the eight gut content samples (sections IV and V) from the initial set of four fish. Of these, 68 representative MAGs (75.0-99.5% complete and 0-6.3% contaminated) were retained for further analysis after cross-sample dereplication (99% ANI threshold) and refinement (Table S2,3), meeting MIMAG medium-high and high quality criteria based on completeness and contamination [37]. Following taxonomic classification most representative MAGs (n = 63) were assigned to a family, while only 43 were further classified to a genus. Approximately 37% of MAGs from the fish gut microbiome therefore belong to previously unidentified genera. MAGs were distributed across eight phyla (Figure 1A). Bacteroidota encompassed the highest number of MAGs (31 all class Bacteroidia), followed by Bacillota_A (n = 23, all class Clostridia) and Bacillota (n = 5, all class Bacilli). The phylum Bacillota is split into multiple groups based on genome phylogeny in GTDB (e.g., Bacillota and Bacillota_A), and represent distinct clades based on our analyses (Figure 1A). The comparatively large number of Bacteroidota, Bacillota_A and Bacillota MAGs recovered in this study is consistent with (a) our previous 16S rRNA gene analysis of 60 *K. sydneyanus* fish luminal samples [10], (b) the unassembled-version of the metagenomic dataset used here [11], which captured more of the rarer community (23% of reads mapped to MAGs in this study) (Table S1), and (c) our analysis of 397 dereplicated MAGs comprising ten fish, which showed no significant differences in relative abundances of the main taxonomic groups (e.g., Bacteroidia, Clostridia and Bacilli, Figure S1A).

**Figure 1.**
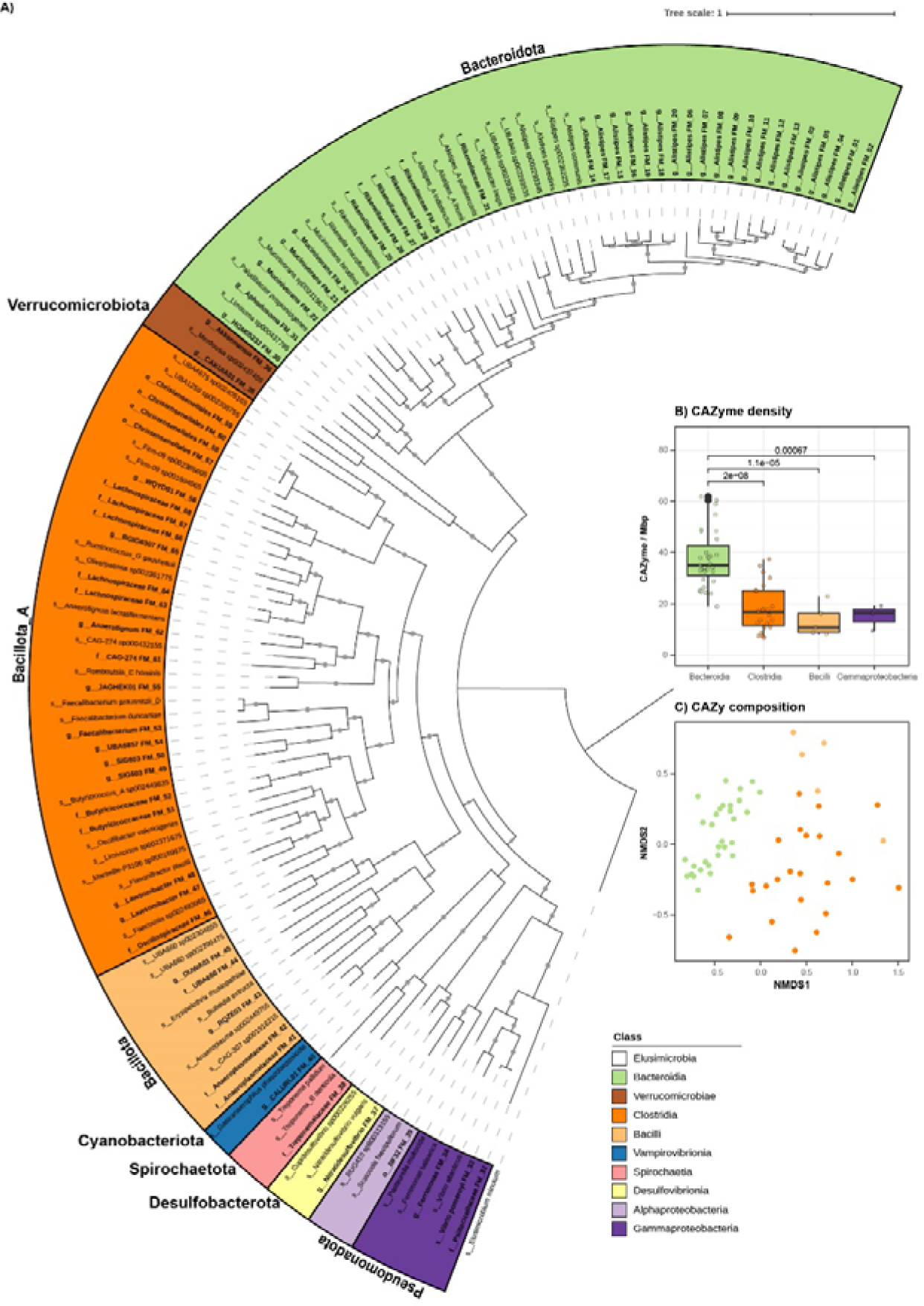
Phylogeny and CAZyme composition of the 68 representative MAGs. (A) Phylogenetic tree of MAGs and references based on the alignment of 120 conserved bacterial markers. Genomes are labelled with their lowest taxonomic resolution classified. Refined MAGs are indicated by a MAG ID and are displayed in bold, and reference genomes are shown unbolded. Clades are coloured by class level and phyla classification by GTDB are indicated by outer labels. Bootstrap values above 70 are shown as solid grey circles. (B) Boxplots displaying the CAZyme density of Bacteroidia, Clostridia, Bacilli, and Gammaproteobacteria classes. Upper brackets indicate the significance (p-values <0.05). The boxes and central line represent the interquartile range and median CAZyme density across MAG in the class, and the whiskers represent the minimum and maximum values within 1.5 times the interquartile range. (C) NMDS of CAZyme composition in Bacteroidia, Clostridia and Bacilli MAGs.

The Bacillota_A comprised species within the families *Lachnospiraceae*, *Oscillospiraceae* and *Acutalibacteraceae* (Figure 1A, Table S3). Bacillota_A is recognized for its fibrolytic capacity in the mammalian gut, in which species of *Oscillospiraceae* and *Lachnospiraceae* are well-established starch and cellulose degraders [38–42]. As butyrate producers, these families are also associated with host health and nutrition [39, 43]. MAGs from Clostridia class Bacillota_A represented the second most CAZyme dense group (Figures 1B, S1B,C), indicating less genetic capacity for glycan metabolism than the Bacteroidota. This is consistent with prior studies showing comparatively fewer CAZymes in the phylum Bacillota than Bacteroidota [44, 45]. Nonetheless, we found that both Bacteroidota and Clostridia contributed substantially more to overall community gene transcription, and CAZyme gene cluster (CGC) transcription than any other taxa group in the hindgut chamber (section V, Figure S2). The hindgut chamber is where the highest microbial biomass and levels of SCFAs occur [12].

The significance of Bacteroidota in complex polysaccharide metabolism is widely acknowledged in the gastrointestinal tract of vertebrates [39, 46, 47], soil [48] and marine environments [30, 49–51]. In these niches, Bacteroidota are key players in carbohydrate biomass recycling due to their large repertoire of CAZymes, CGCs, and transporters dedicated to the saccharification of specific complex carbohydrates [39, 49, 51, 52]. In this study, Bacteroidia exhibited a significantly higher density of CAZymes compared to other taxonomic classes (Figures 1B). This observation was consistent across the gut communities of fish collected in 2017 and 2020 (Figure S1B,C). Bacteroidia also exhibited a distinct CAZyme composition (Figure 1C), implying that these Bacteroidia have a distinct carbohydrate utilization capacity. Among Bacteroidia MAGs, the genus *Alistipes* represented the largest taxonomic group (n = 20/31 MAGs), and formed a distinct clade from other Bacteroidia and *Alistipes* from other habitats. The closest cultivated relatives to this clade were *Alistipes communis* [53] and *Alistipes putredinis* [54], both of which are primarily described from human clinical studies [54] (72.2-75.7% ANI shared with 16 *Alistipes* MAGs, Table S4).

Increases in *Alistipes* relative abundance in the human gut have been attributed to animal protein-rich diet treatments (e.g., eggs, bacon, pork and beef) [55], implying the involvement of this genus in protein degradation. In mice, plant-based diets (e.g., comprising resistant maltodextrin, fructo-oligosaccharide, galacto-oligosaccharides, iso-malto-oligosaccharides) are associated with a decline in the relative abundance of *Alistipes* [56, 57]. In contrast, metagenomic rumen studies imply *Alistipes* have the capacity to utilize oligosaccharides from plant-cell walls. For example, *Alistipes* increased in relative abundance in rumen metagenomes of Holstein cows fed on a high-forage (HF) diet, and also displayed higher abundances of glycoside hydrolases (GHs) and carbohydrate binding modules (CBMs) [58]. The predominance of *Alistipes* within the large intestine of ruminants has also been attributed to the utilization of host glycan, based on high numbers of CAZy families GH109, GH20 and GH92 encoded by these organisms [59]. While the role of *Alistipes* in various hosts remains under-explored, studies collectively suggest that species within this genus perform differing host-dependent roles related to substrate degradation [54]. *Alistipes* in *K. sydneyanus* are thus predicted to degrade the polysaccharides of seaweeds in the host diet, and their potential to do so, alongside other carbohydrate degraders, is illustrated below.

### Bacteroidia and Clostridia species are capable of initiating polysaccharide degradation

Genomic analyses revealed that the microbial community in the hindgut of *K. sydneyanus* encoded a diverse array of enzymes to attack complex carbohydrates in the fish diet (Figure 2), and completely degrade these to monosaccharides (e.g., glucose, galactose, fucose and uronic acids). Nevertheless, enzymes acting on high molecular weight (HMW) substrates, such as guluronate-specific alginate lyases, glucan endo-1,6-β-glucosidases, glucan endo-1,3-β-D-glucosidases, endo−1,3(4)−β−glucanases, α-fucosidases, κ-, ι-, λ-carrageenases and β−agarases were found primarily within a small number of Bacteroidia and Clostridia MAGs. The capacity for initiating the hydrolysis of HMW polysaccharides into smaller polymers for membrane transport is a critical step in enabling the utilization of dietary substrates [60]. Altogether, the substantial representation of Bacteroidia and Clostridia within the gut communities, alongside their endo-acting enzymatic capacity, suggest that they play a pivotal role in the degradation of substrates derived from the fish diet. This observation is in line with the current status of Bacteroidota as efficient polysaccharide degraders in diverse environments [45, 50, 61–63]. However, this result also suggests the microbial drivers of *K. sydneyanus* seaweed degradation are somewhat unique. Studies on other herbivorous fish species, such as *Acanthurus sohal*, *Naso elegans* and *Naso unicornis*, attribute the cleavage of dietary polysaccharides primarily to Bacillota_A encoded endo-β-1,4-glucanases (GH74), β-agarases (GH50 and GH86), and β-porphyranases / κ-carrageenases (GH16) [64].

**Figure 2.**
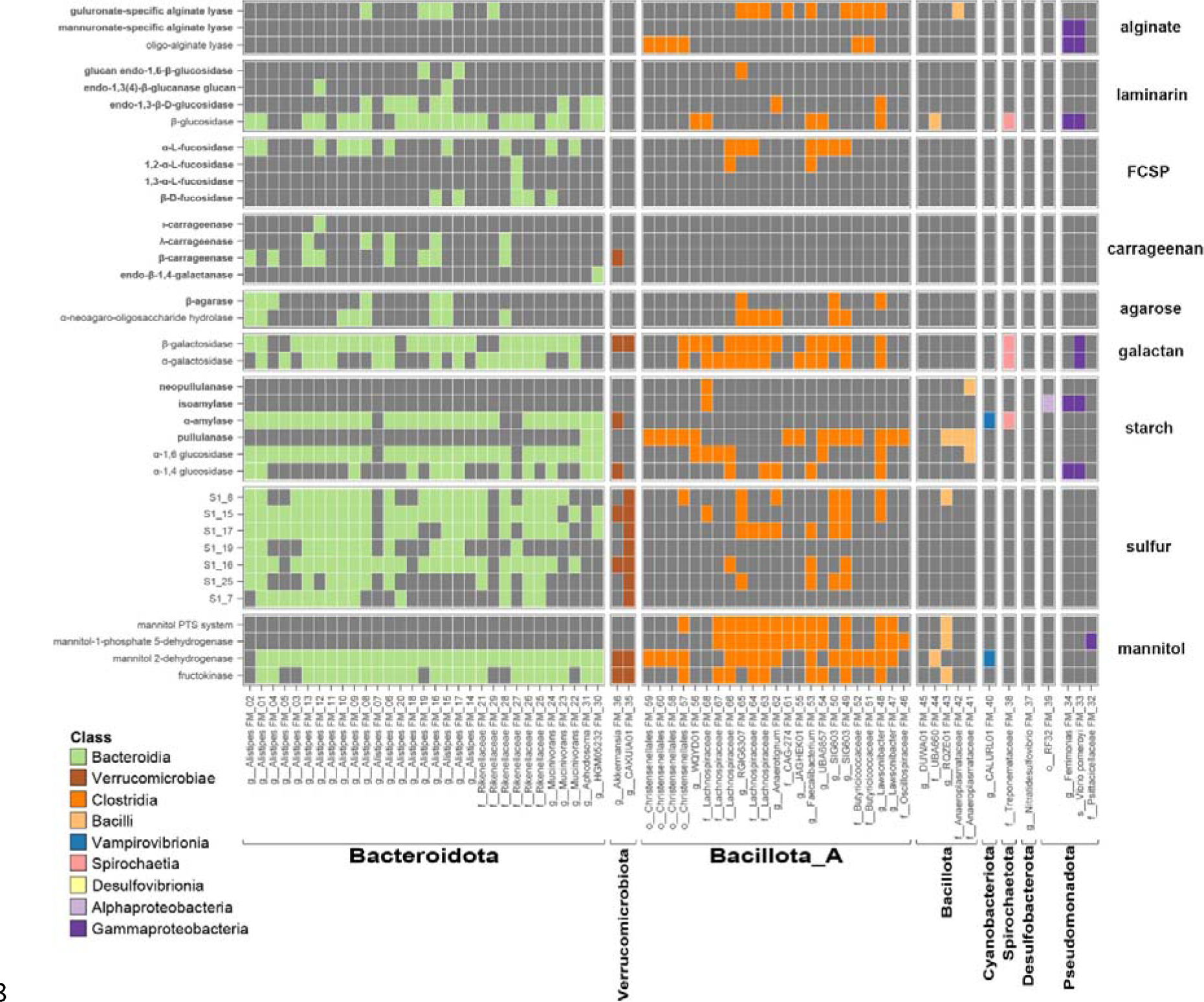
Heat map showing the presence of genes associated with carbohydrate degradation in each MAG. Gene presence is indicated by boxes coloured by taxonomic assignment at the class level, and absence is indicated by grey shading. MAGs, shown along the x-axis, are ordered by their phylogenetic placement in Figure 1a. Labels at the bottom in bold indicate Phyla. Predicted enzymes are denoted on the y-axis on the left, and predicted substrates are shown on the right. Predicted CAZymes known to act on HMW polymers (based on their EC number) are shown in bold. The upper seven plot panels display the presence of CAZymes targeting the main substrates of brown and red algae (panels alginate to starch). The eighth panel displays the presence of sulfatases associated with FCSP and carrageenan desulfation. The bottommost panel shows the presence of genes associated with mannitol utilization.

Encoded enzymes targeting low molecular weight (LMW) substrates such as β-glucosidases, β-galactosidases and α-galactosidases were widely distributed across genomes within Bacteroidia and Clostridia (Figure 2). Of these, β-glucosidase genes are associated with the utilization of oligosaccharides derived from laminarin, while both β-and α-galactosidases are implicated in the downstream degradation of galactose-rich polysaccharides such as FCSP, carrageenan and agarose. These are the degradation products of substrates acted on by predicted HMW targeting enzymes encoded by members of the same microbial community. Hence a larger number of MAGs containing only LMW acting capacities likely depended on the extracellular hydrolyzation of HMW substrates undertaken by a small number of keystone Bacteroidia and Clostridia in order to grow [65].

Encoded enzymes associated with the hydrolysis of starch, including HMW (e.g., neopullulanases, isoamylases, α-amylases and pullulanases) and LMW forms (e.g., α-1,6 glucosidases and α-1,4 glucosidases), were both widely distributed in the community (Figure 2). However, the utilization of starch is unlikely to be a major component of adult *K. sydneyanus* microbial metabolism, as starch constitutes less than 3% (dry weight) of the nutritional composition of dietary red algae such as *Gigartina livida* [5]. Also, the effect of acid lysis in the *K. sydneyanus* stomach, in addition to the high activity levels of amylases found in the anterior intestine (>300 µg reducing sugar ml^-1^ min^-1^ in section II), may digest starch completely or leave very little amounts of starch in later portions of the gut [5, 6].

Polysaccharides such as FCSP and carrageenan are heavily associated with sulfate esters (up to 40% of its dry weight) [66–68]. To consume these polysaccharides, the sulfate groups in the sugar backbone require removal prior to hydrolysis [67, 69]. Released sulfate can then be utilized as an energy source by sulfate-reducing bacteria or converted into sulfur-containing amino acids by the community [70, 71]. In the present study, Bacteroidia MAGs encoded an enriched and expanded repertoire of sulfatase families associated with FCSP and carrageenan desulfation (Figure 2, Table S7). These functions were scarce in genomes from Clostridia. Among the sulfatase families identified, S1_15, S1_16 and S1_17 are commonly associated with sulfate removal from FCSP derived from *Fucus vesiculosus* (sulfation on positions C2 and C3) or *Cladosiphon okamuranus* (sulfation on position C4) [67]. The sulfatase family S1_25 is an exo-sulfatase acting on position С3 of fucose [72]. The ι- and κ-carrageenan isoforms depend on sulfatase families S1_19 or S1_7 to remove sulfate esters on position C4 of its galactose blocks [73]. Additionally, ι-carrageenan requires prior removal of sulfate esters on position C2 of 3,6-anhydro-D-galactose moieties by sulfatase family S1_17 [73, 74]. Lastly, the sulfatase family S1_8 is implied to perform the desulfation on position C2 of the galactose moieties of λ-carrageenan [75]. Altogether, the enrichment of genes necessary for HMW sulfated polysaccharide depolymerization further emphasises the key role predicted for Bacteroidota in initiating the deconstruction of polysaccharides from the fish diet.

The ability of the microbiota to ferment mannitol likely underpins the complete depletion of this sugar-alcohol, as observed along the hindgut gradient of *K. sydneyanus* [76]. This sugar-alcohol could support the growth of microbes with a limited capacity to consume complex carbohydrates in early sections of the fish gut [76]. We found that genes for mannitol utilization via mannitol 2-dehydrogenase (M2DH) and fructokinase were mainly present among Bacteroidia, Clostridia and Verrucomicrobiae MAGs (Figure 2) enabling conversion of D-mannitol to phosphorylated fructose (D-fructose-6P) in the glycolysis pathway. However, only the Clostridia encoded mannitol PTS channel and mannitol-1-phosphate 5-dehydrogenase (M1PDH) genes [77, 78], facilitating another route from D-mannitol to D-fructose-6P. The presence of both pathways for mannitol utilisation may confer a competitive advantage to Clostridia for mannitol usage.

### The *Alistipes* are specialized degraders of polysaccharides

Investigation of carbohydrate-acting gene arrangements can provide valuable information for elucidating the degradation pathways for specific glycans and discovering novel CAZyme families [79]. CAZymes encoded by the same CAZyme Gene Cluster (CGC) generally feature a set of complementary enzymes capable of fully deconstructing a specific glycan (e.g., a glucan endo-1,6-β-glucosidase and β-glucosidase to hydrolyse laminarin into glucose), and as such facilitate glycan degradation via their co-expression [27, 51, 65]. Analysis of the *K. sydneyanus* gut community revealed a total of 1257 CGCs, in which 283 contained at least two degradative CAZymes (e.g., GH and/or PL and/or CE) (Figure 3A,B). Among these degradative CGCs, 135 could be linked to substrates relevant to the fish diet, such as alginate, laminarin, FCSP, carrageenan, agarose and starch (Figure 4A). Accordingly, these CGCs often encoded enzymes such as alginate lyases (PL17, PL34 or PL6), laminarinases (GH30, GH149 or GH16), fucosidases (GH29, GH30 or GH141), carrageenases (GH150), agarases (GH16 or 86) and amylases (GH13 or GH133 or GH57 or GH77) (Table S9).

**Figure 3.**
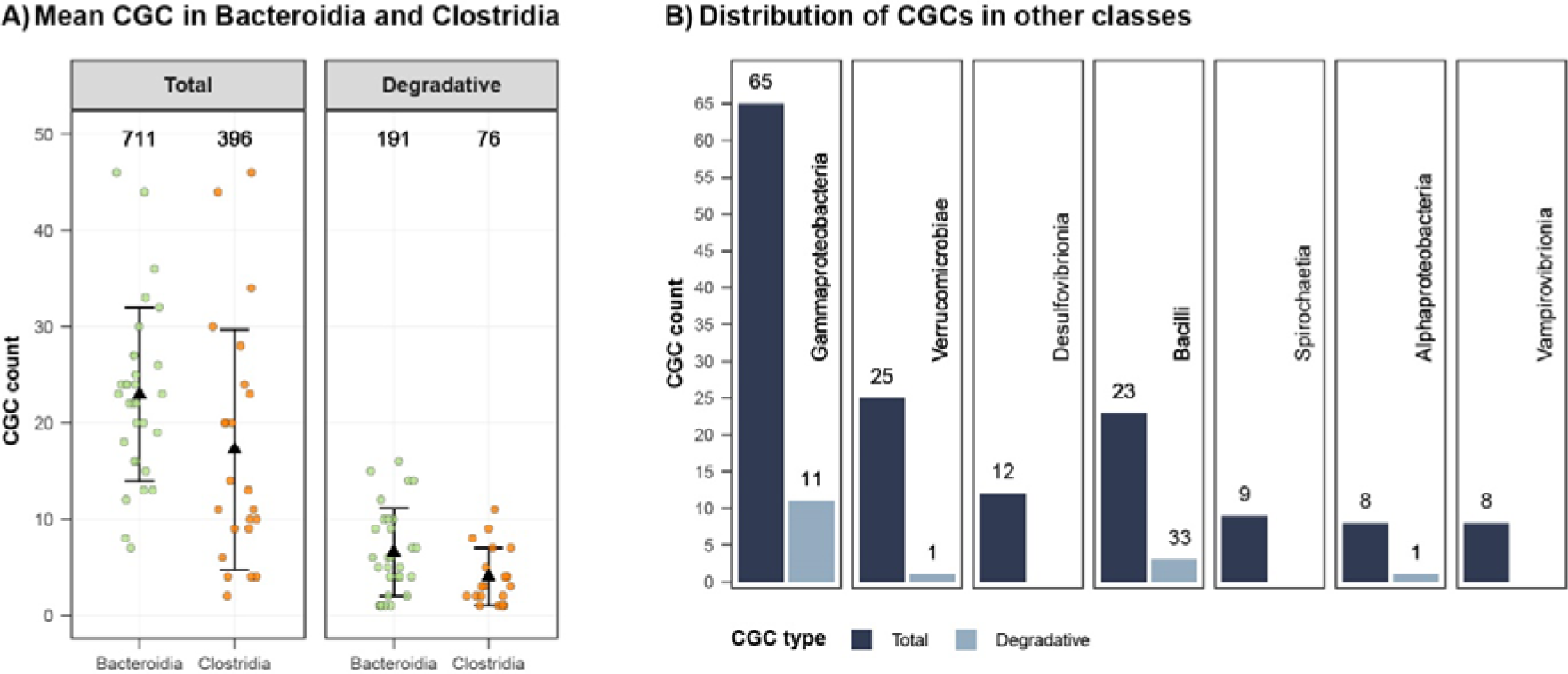
Summary of CGC types across class and their degradative CAZyme composition. (A) Strip plots showing the total and degradative number of CGCs encoded per MAG in Bacteroidia (green) and Clostridia (orange). Degradative CGCs are defined here as those containing at least two degradative CAZymes. Black triangles indicate the mean of CGCs per class (across MAGs) and error bars show the standard deviation. Numbers at the top display the sum of each CGC type per taxonomic class. (B) Bar plots indicating the sum of total and degradative CGCs in other classes, which each have less than 3 MAGs containing degradative CGCs. Numbers above bars show the CGC count.

**Figure 4.**
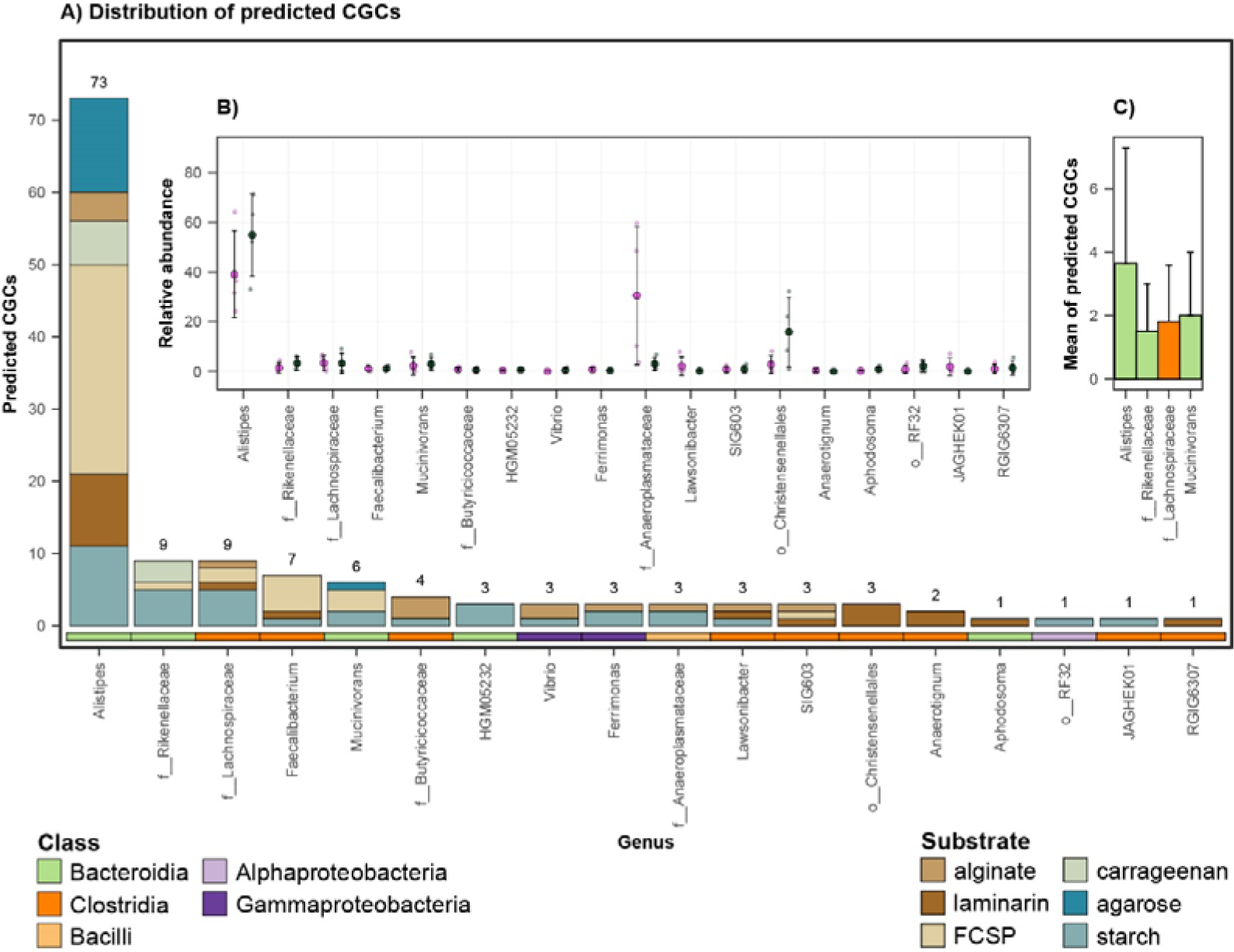
Classified CGCs with degradative properties associated with substrates common to the fish diet. (A) Number of predicted CGCs per genus. Stacked bars are coloured according to their predicted substrate. Tiles below bars indicate taxa class affiliations. (B) Relative abundances in sections IV and V of genera containing predicted CGCs. Small open circles represent the genus relative abundance in each fish. Mean relative abundance values are shown by large circles coloured by sections (IV = pink, V = dark green), and error bars denote standard deviations. (C) Bar plot of the average number of predicted CGCs per MAG in the top four genera with more than two MAGs. Bars are coloured by class affiliation and error bars indicate the standard deviation.

Inspection of substrate-assigned CGC distributions across the gut communities (associated with the *K. sydneyanus* diet) revealed a sizeable and abundant collection of CGCs within the genus *Alistipes* (n = 73, across 20 MAGs) (Figure 4A). Per MAG, the average number of CGCs in *Alistipes* was 3.65 ± 3.34 S.D. (followed by *Mucinivorans* with 2 ± 1 S.D. algae-associated CGCs per MAG) (Figure 4C, Table S11). Results imply both redundancy and diversity in the carbohydrate utilization pathways employed by *Alistipes*, considering the distribution of genes with CAZyme-encoding functions across the group (guluronate-specific alginate lyases = 6, glucan endo-1,3-β-D-glucosidases = 13, α-L-fucosidases = 79, κ-carrageenases = 8 and λ-carrageenases = 5) (Table S10). In addition to CGC diversity in *Alistipes*, the genus displayed considerable variation in relative abundance across fish in both hindgut sections sampled (section IV = 39.1% ± 17.49 S.D. and section V = 54.92% ± 16.48 S.D.), potentially resulting in variable metabolic significance across fish replicates (Figure 4B). The higher average relative abundance of *Alistipes* in section V (albeit not a statistically significant difference, Kruskall-Wallis p-value = 0.25) (Table S11), may also reflect different stages of glycan utilisation along the fish gut. These results are supported by our previous 16S rRNA gene analysis of 60 *K. sydneyanus* fish showing that the genus *Alistipes* was significantly more abundant in section V than in sections III and IV [10].

### Evidence for functional redundancy and niche partitioning among co-abundant taxa

The utilization of glycans is a significant driver of interactions yielding positive and negative synergies within microbial gut communities [41, 80–82]. Moreover, the wide range of non-utilized glycans by the *K. sydneyanus* host [6], and long residence time of digesta [83], support a landscape for complex inter-species interactions through resource coworking or competition [82]. We therefore examined pairwise correlations of MAG relative abundances to elucidate the potential for positive and negative synergistic relationships based on taxa co-abundance (Figure 5). A clear delineation was observed among co-abundant MAGs when considering changes in their relative abundance from gut section IV to V. Despite a clear trade-off between Clostridia and Bacteroidia relative abundances, and transcriptional activity, between these sections (Figures 4B, S2), all six co-abundant groups of taxa (A1 to B4) displayed a diverse assortment of taxonomic affiliations, highlighting the potential for inter-species collaborations (or competition). In particular, the presence of CGC-rich *Alistipes* MAGs, across multiple co-abundant groups (Figure 5), provides further evidence that this genus is important for carbohydrate metabolism within the *K. sydneyanus* gut.

**Figure 5.**
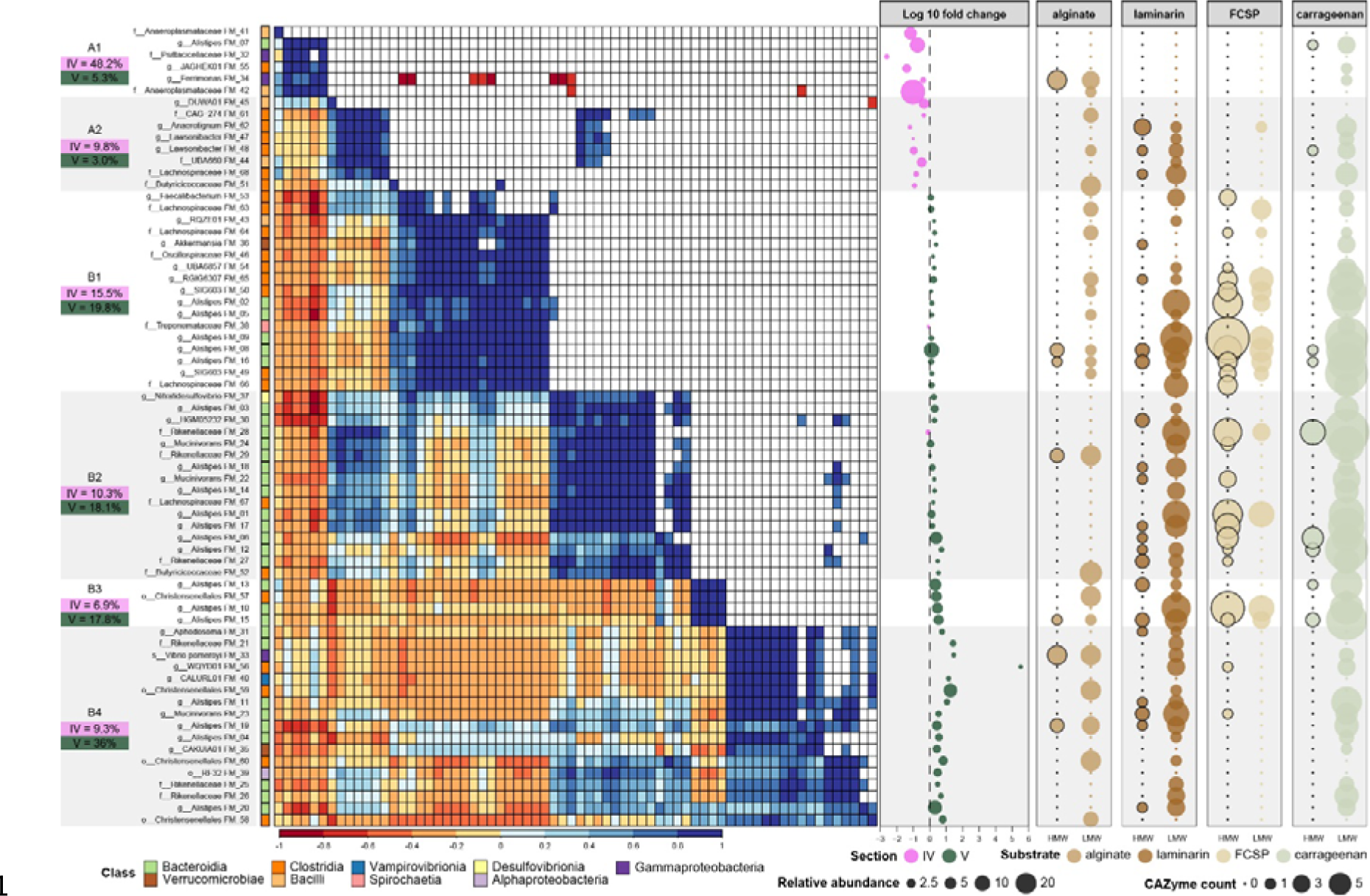
MAG co-abundances and CAZyme enrichment across co-abundant taxon groups. Leftmost plot: Pearson correlation heatmap of MAG relative abundances indicating positive (blue shading) and negative (red shading) correlations (lower triangle). White boxes in the upper triangle denote insignificant p-values (> 0.05). Leftmost labels delineate co-abundant groups and display their group summed relative abundance in sections IV (pink) and V (dark green). Labels immediately left of the heatmap give MAG IDs and their class affiliation (outer coloured tiles). Middle plot: Log10 fold changes in the relative abundance of each MAG between gut sections. Bubble sizes and colours represent relative abundance and the section the MAG was most abundant. The four right hand side panels display the CAZyme enrichment in each MAG associated with the utilization of high molecular weight (HMW) and low molecular weight (LMW) isoforms of the target polysaccharides.

We observed an ample distribution of CAZyme-encoding genes for β-glucosidases (laminarin LMW), galactosidases (FCSP/carrageenan LMW) and oligo-alginate lyases (alginate LMW) across co-abundant groups, supplied by numerous taxa (Figure 5). Results therefore indicate a high level of functional redundancy in the gut communities based on the taxonomically widespread and relatively abundant exo-acting CAZyme capacity. Such redundancy likely confers the microbial community with flexibility in handling a diverse diet [82, 84], and may be facilitated by niche partitioning (e.g., based on variations in enzyme efficiencies or substrate prioritization depending on the nutrient landscape) [80, 82].

The group-wise changes in relative abundance from gut section IV to V presumably reflects a sequential breakdown of glycans along gut sections by taxa in the A groups (exclusively more abundant in section IV) and B groups (members abundant equally in sections IV and V or enriched in V) (Figure 5, S3A). The emergence of B groups along the gut implies the start and progression of intensive glycan metabolism in gut section IV, which is supported by their large and diverse combined CAZyme capacity (Figure S3). This coincides with the significantly higher levels of total SCFA in *Kyphosus sydneyanus* gut sections IV and V compared to section III (previously reported total SCFA concentrations were 19.35 ± 4.3 S.E. in section III, 49.13 ± 2.62 S.E. in section IV, and 58.07 ± 4.62 S.E. in section V) [12].

The A groups, enriched in section IV only, encoded a limited glycan degradative capacity compared to B (Figure 6). This observation, combined with the sharp decline in relative abundance of the two A groups from section IV to V, indicates that they are outcompeted in section IV (Figures 6, S3B). Nonetheless, the high relative abundance in section IV of group A1 taxa, Bacilli (FM_42), which encodes an oligo alginate lyase, and Bacteroidia (FM_07), which has one κ-carrageenase and four β-galactosidases genes, suggest that alginate and carrageenan metabolism could commence in section III before increasing in later sections. Previous work demonstrated low enzymatic activity against alginate and κ-carrageenan (≤ 1 µg reducing sugar ml^-1^ min^-1^) between sections II and III [6]. Group A2 MAGs displayed a slightly broader carbohydrate capacity attributed primarily to Clostridia, potentially enabling the utilization of alginate oligosaccharides, laminarin and carrageenan (Figure 5,6). MAGs in this group also displayed moderately positive correlations in relative abundance with subsets of MAGs from groups B2 and B4, which share a common capacity for laminarin and carrageenan degradation (via α- and β-desulfated galactans) (Figure 5).

**Figure 6.**
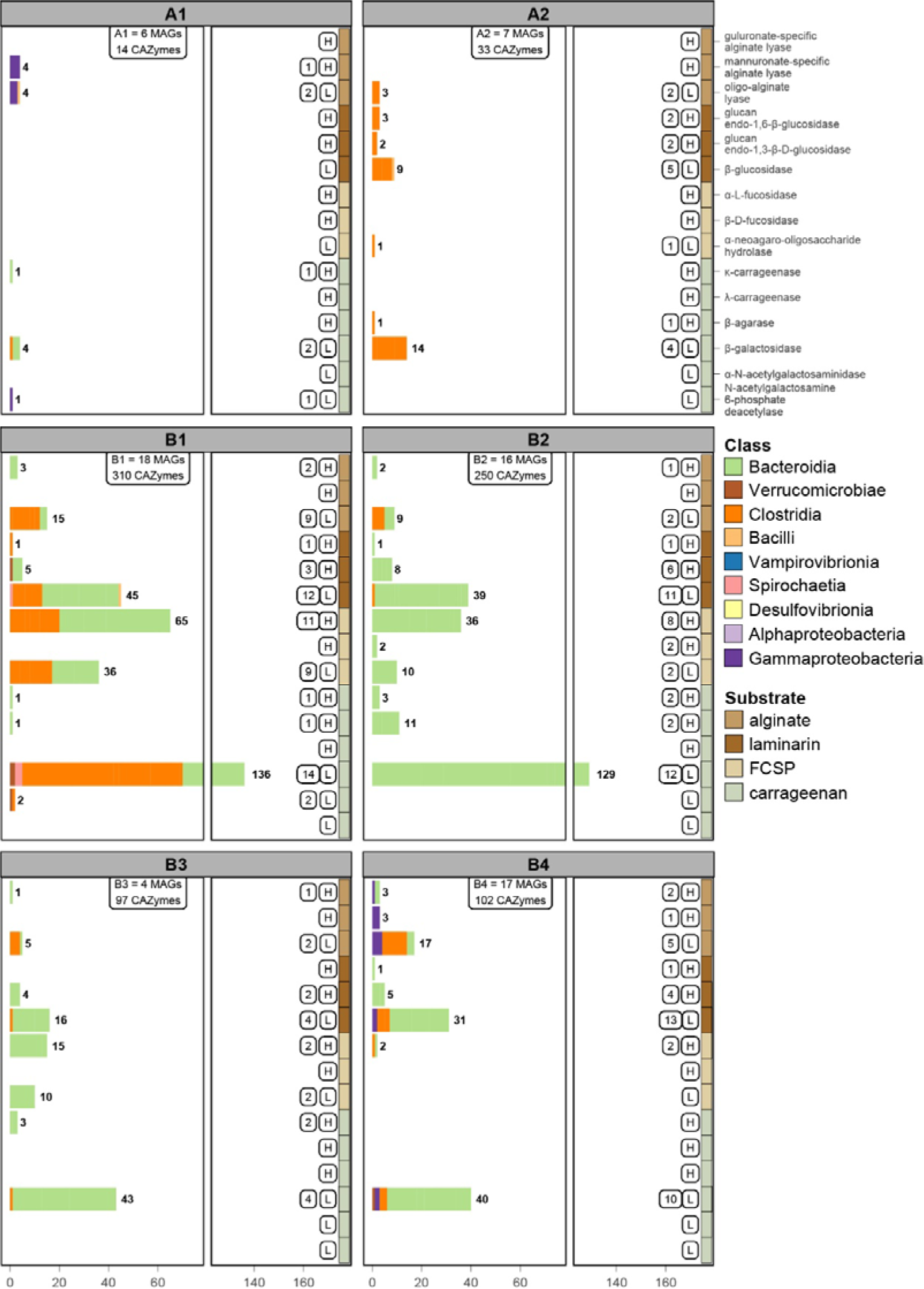
Stacked bar plots showing the pooled count of genes in each co-abundant taxa group per enzyme encoded. X-axis = gene counts (also shown numerically besides bars). Bars are coloured according to taxon class. A break was inserted in the x-axis to accommodate the large number of galactosidases encoded by Bacteroidia MAGs in groups B1 and B2. Coloured tiles on the right represent the substrate each enzyme is predicted to act on. Boxes besides the coloured tiles indicate the number of MAGs encoding specific enzymes, and their capacity for HMW (H) or LMW (L) substrates.

The four B groups encode a diversified carbohydrate degradation capacity, with a large enrichment in encoded enzymes, such as β-galactosidases, α-fucosidases and β-glucosidases (Figures 5, 6). Their rich CAZyme repertoires explain their maintained or increased relative abundance between gut sections IV and V. Despite the broadly analogous carbohydrate degrading potential among B groups, B1 demonstrated the widespread presence of CAZymes within Clostridia MAGs that are associated with the breakdown of alginate, laminarin and FCSP. In contrast, group B2 comprised mainly Bacteroidia, including three MAGs with the highest number of carrageenase genes (for HMW carrageenan) across all co-abundant groups (Figures 5,6). This may have provided an advantage to B2 members in sourcing galactose from carrageenan in addition to FCSP. On the other hand, only two B2 MAGs carried alginate lyases, suggesting little preference of this group for alginate (Figure 5). The large proportion of Bacteroidota, including seven *Alistipes* MAGs, within B2 implies a metabolism driven by CGCs given the substantial enrichment of CGCs in the genus *Alistipes* (Figure 3). Members of group B2 may thus have benefited from the adaptive regulation afforded by CGCs to rapidly changing nutrient conditions [46].

Groups B3 and B4 exhibited a tendency for greater relative abundance in hindgut section V. Group B3 consisted of only four MAGs comprising *Alistipes* (n = 3) and Christensenellales (n = 1). These MAGs encoded a relatively large number CAZyme genes (n = 97), despite there being far fewer MAGs in this group (CAZyme encoding genes were present at a frequency of 24 per genome in group B3, and 6 to 17 per genome in the other B groups) (Figure 6). However, group B3 lacked the capacity to produce some key enzymes, including glucan endo-1,6-β-glucosidases and mannuronate-specific alginate lyases that are involved in the initial depolymerization of laminarin and alginate, respectively [85]. As a result, bacteria in this group may not efficiently source nutrients from these glycans on their own, and potentially required an initial hydrolyzation of laminarin and alginate to thrive. This may explain their increase in abundance in section V, where depolymerized forms of laminarin and alginate are present as a result of previous laminarinase and/or alginate lyase activity [6], presumedly from groups A1 or A2. In addition, Group B3 could be supported by substrates derived from other B groups encoding mannuronate-specific alginate lyase (B4) or glucan endo-1,6-β-glucosidases (B1, B2 and B4) (Figure 6). Overall, the carbohydrate degrading capacity in B3 and B4 suggested distributed capacity for LMW substrates across these groups. While B4 exhibited greater capacity towards HMW substrates, only B3 possessed carrageenases, and B4 harboured substantially fewer fucosidases, indicating a deficiency in utilising galactose on its own (Figure 5, 6).

These results highlight the widespread presence and abundance of genes encoding enzymes targeting LMW carbohydrates across taxa in both gut sections (610 LMW targeting genes across 60 MAGs), enabling utilisation of the breakdown products from a smaller group taxon capable of breaking down HMW substrates (184 HMW targeting genes across 38 MAGs) (Tables S10). Despite the redundancy observed across the gut community, nuances in CAZyme encoding may confer competitive advantages to each taxon group, depending on the nutrient landscape, that regulate the relative abundance of overlapping groups, or their presence-absence, along the intestinal tract of the fish (Figure S3A,B). These overlapping groups are likely able to coexist by utilizing alternative nutrient sources or prioritizing substrates to avoid direct competition [46, 80, 82].

## CONCLUSIONS

Our results demonstrate a large capacity for microbial carbohydrate degradation along the *K. sydneyanus* gut associated with the fish diet of brown and red macroalgae. Bacteroidia (including many novel *Alistipes*) and Clostridia MAGs contained the highest CAZyme-encoding gene densities in the community. These phyla were also predicted to encode endo-acting CAZymes essential for the initiation of glycan depolymerization. Of these, Bacteroidia MAGs encoded an expanded repertoire of sulfatases, indicating an enhanced capacity to utilize sulfated polysaccharides such as FCSP and carrageenan in the fish diet. The *Alistipes* contained a remarkable diversity of CGCs, suggesting a specialization for carbohydrate utilization and adaptive regulation in changing nutrient conditions. Analysis of MAG co-abundance dynamics between gut sections IV and V revealed multiple cohorts with varying abundances capable of degrading dietary carbohydrates. The higher prevalence of Clostridia in groups more abundant in the proximal gut section indicates their role initiating polysaccharide utilization prior to Bacteroidia, and is consistent with their relatively high transcriptional activity in section IV. Diverse Clostridia, in addition to Bacteroidia, were also present in groups abundant in section V. Among these taxa, a ubiquity of encoded galactosidases and β-glucosidases indicates their intrinsic ability to compete for resources, and substantial functional redundancy in CAZyme-encoding in the fish gut. The range of CAZymes encoded per group also indicates high versatility in glycan utilization, and thus the capacity for niche differentiation. Accordingly, results indicate the potential for inter-species collaborations based on the diverse assortment of taxa that co-occurred with complementary carbohydrate degradation potentials. Multiple Clostridia and Bacteroidia taxa can thus contribute to the degradation of *E. radiata*, *C. maschalocarpum*, *G. macrocarpa* and *C. ustulatus* in the *K. sydneyanus* hindgut, and are of importance for host nutrition.

## ACKNOWLEDGEMENTS

We thank Alessandro Pisaniello, Bikiran Pardesi, Brady Doak and Peter Browne for assistance with field and laboratory work. Computational resources were provided by New Zealand eScience Infrastructure. The study was funded by the New Zealand Ministry of Business, Innovation and Employment Endeavour Grant (project UOAX1808-CR-2) awarded to K.D.C. and W.L.W.

## DATA AVAILABILITY

Metagenomic data was deposited with NCBI under BioProject PRJNA1029302 (MAG accession numbers are given in Table S3).

## COMPETING INTEREST

The authors declare that they have no competing interests.

## Notes

### Competing Interest Statement

The authors have declared no competing interest.

