## Supplementary Information for "Bacteroidia and Clostridia genomes collectively encode for a progressive cascade of marine polysaccharide degradation along the hindgut of the herbivorous fish *Kyphosus sydneyanus*"

**SUPPLEMENTARY METHODS**

**Metagenome data generation from the 2017 fish collection**

Lumen samples from the four individuals collected in 2017 (approval 001636 from the University of Auckland Animal Ethics Committee) were extracted using the PowerSoil® DNA Extraction Kit (Mo Bio Industries, Ltd., Carlsbad, USA) and purified using the Agencourt AMPure XP kit (Beckman Coulter, Brea, California, United States) [1]. Library preparation and sequencing was undertaken by Auckland Genomics (University of Auckland, Auckland, New Zealand) using the NexSeq® AmpFREE Low DNA Library Kits (Lucigen, Middleton, USA) and the Illumina NextSeq 500 platform (Hayward, CA, United States), generating 131.6 Gb of sequence (2 × 150 bp reads.) (Table S1) [1].

Adapters were removed from whole genome shotgun sequencing reads using cutadapt version 2.3 [2] (see Supplementary Information for trimming and MAG generation of the 2020 samples). Sequences were then trimmed using Sickle version 1.33 based on a minimum quality score of 25, retained if ≥ 80 bp [3], and evaluated using FastQC version 0.11.7 [4]. Trimmed reads from each sample were assembled separately using SPAdes version 3.11.1 with –meta -m 100 -t 16 -k 43, 55, 77, 99, 121 parameters [5]. To determine differential coverage for genome binning, trimmed reads were mapped to contigs using Bowtie version 1.2.0 with the following parameters: --phred 30 -quals -n 1 -l 222 --minins 200 --maxins 800 –best [6]. Contigs were binned using MetaBAT version 2.12.1 [7], MaxBin version 2.2.4 [8], and CONCOCT version 0.4.1 [9] utilizing differential coverage and tetranucleotide frequencies. The resulting bins were compared, and representatives were selected using DAS Tool version 1.1.1 [10]. To generate a set of unique representative MAGs, genomes were dereplicated across samples using dRep version 1.4.3 with the default 99% average nucleotide identity (ANI) threshold [11]. MAG refinement was performed using VizBin incorporating differential coverage [12]. CheckM version 1.0.12 was used to estimate refined MAG completeness and contamination [13]. Mapped read counts were obtained using the BBTools ‘pileup.sh’ script [14] from Bowtie sam files [6]. Counts were normalized by library size and genome length to estimate the genome coverage (and hence relative abundance) of MAGs across samples.

**Subsequent 2020 fish collection and omics data generation**

Six *K. sydneyanus* individuals were collected from waters around Great Barrier Island, Auckland, New Zealand, in January 2020 (approval 001949 from the University of Auckland Animal Ethics Committee). Gut contents from sections III, IV and V were homogenized, aliquoted into microtubes and immediately stored in liquid nitrogen until storage in a -80°C freezer. Prior to extraction, samples were thawed and centrifuged at 15000 x g for 2 min and the supernatant removed. The leftover pellet was reconstituted using solution CD1 from the PowerSoil Pro kit (QIAGEN, Germantown, MD, United States), and DNA was extracted following manufacturer instructions. Extracted DNA was purified using Genomic DNA Clean & Concentrator-10 (Zymo Research, Irvine, California, United States).

DNA libraries were prepared at the Otago Genomics Facility (University of Otago, Dunedin, New Zealand) with the Takara Thruplex DNA-Seq 96D kit (Takara Bio, Kusatsu, Shiga, Japan) for 18 high molecular DNA samples from sections III, IV and V of the six fish. Paired 125 bp reads were generated using the Illumina HiSeq V4 platform (Illumina, San Diego, CA, United States).

RNA was extracted with the Monarch^®^ Total RNA Miniprep Kit (New England BioLabs, Ipswich, Massachusetts, United States) following manufacturer instructions (including a DNA removal step). RNA yields were measured using Qubit and purity using a NanoPhotometer. DNA removal was confirmed by no amplification of the 16S rDNA gene in extracted versus positive control samples [15]. RNA was purified using the RNA clean and concentrator-5 kit (Zymo Research, Irvine, California, United States) followed by the AMPure XP kit (Beckman Coulter, Brea, California, United States) to improve 260/230 purity ratios. Integrity of RNA was then evaluated using a 2100 Bioanalyzer and RNA Analysis kit (Agilent, Santa Clara, California, United States), Libraries were prepared for a subset of 13 samples comprising three section III, four section IV and six section V (showing RIN values from 5.5 to 7.6) using the Zymo-Seq RiboFree® Total RNA Library Kit (Zymo Research, Irvine, California, United States) by Otago Genomics. Sequencing was performed using three Illumina NextSeq 2000 P3-200 flow cells outputting a minimum of 110 Gb per sample of 2 x 100bp PE reads.

Raw metagenomic reads were trimmed and adapters removed using BBMap version 37.93 with the BBDuk tool (parameters ktrim=r, k=35, mink=20, hdist=1, tpe, tbo, qtrim=rl, trimq=30, minlen=80) [14]. Assembly was performed per sample using SPAdes version 3.15.4 [5] (parameters --meta -m 100 -t 16 -k 43,55,77,99,121). Binning was performed using MetaBAT version 2.13 [7], MaxBin version 2.2.6 [8] and CONCOCT version 1.1.0 [9]. Contig coverages used in the binning process were generated using Bowtie version 2.3.5 [6] (parameters --phred33 -p 12 -N 1 -L 32 --minins 200 –sensitive --maxins 800), mapping each sample read set to their respective assembly. DAS_Tool version 1.1.1 [10] was used to keep the best metagenome-assembled genomes (MAGs) per binning tool per sample, resulting in 448 MAGs.

To compare 2017 and 2020 datasets, the newly generated MAGs from six *K. sydneyanus* individuals were combined with 197 generated in this study (Table S2). All MAGs were checked for completeness and contamination using CheckM version 1.2.1 [13]. Only genomes >70% complete and with <5% contamination were kept. These genomes were then dereplicated using dRep version 2.3.2 [11] using the default threshold of 99% average nucleotide identity (ANI) to create a representative set of genomes. These were classified using the GTDB Toolkit (GTDB-Tk, version 2.1.0) against the release 214 database [16]. The final dataset contained 397 unique good quality *K. sydneyanus-*associated genomes.

RNA reads were trimmed using trimmomatic version 0.39 [17] with adapter removal (trimmomatic parameters HEADCROP:10 SLIDINGWINDOW:4:30 MINLEN:70). The rRNA reads were removed using SortMeRNA version 2.1 [18]. Non-rRNA reads were mapped to these 397 unique good quality genomes and their counts normalised by gene length and library size (number of reads mapped to gene)*(1000/gene length)*(1000000/library size)).

**Substrates estimation based on EC similarity**

Genes annotated as CAZymes with similar EC functions were filtered based on their association to paths of degradation of brown and red algae polysaccharides. Each EC function was assigned a substrate according to their CAZy annotation and the literature. EC functions lacking characterization such as “EC 4.2.2.-“ and “EC 3.2.1.-“ were assigned a substrate if the corresponding EC was the only uncharacterized activity within the respective the CAZy family (e.g., PL6 contained only one “EC 4.2.2.-“). CAZy families that contained redundant functions, such as GH110 (α-galactosidase and α-1,3-galactosidase) or accommodated unique EC functions (e.g., GH82: ι-carrageenase 3.2.1.157, or GH150: λ-carrageenase 3.2.1.-) were also assigned the EC the unique function regardless of presenting similarity with any EC characterized CAZyme. Substrate estimations and ECs are detailed in Table S5.

**Mannitol genes identification**

An initial keyword search across UNIPROT, Uniref100, KEGG and Pfam annotations was performed to identify genes associated with mannitol utilization pathways [1]. Next, these genes were inspected using a five gene sliding window to identify potential mannitol operons containing a minimum of two genes, in which, at least one gene contained the keyword “mannitol” in its annotation. A last filter considered their protein domain annotations to estimate their probable gene function. The keywords used in this search are detailed in the Table S6.

**CGC classification and substrate assignment**

In order to assign CGCs to a substrate, only CGCs containing two degradative (GH, PL or CE) CAZy families were considered. These CGCs were inspected manually considering the CAZyme content of the CGC, their similarity with EC functions, and their CAZy annotation. A substrate was assigned if the arrangement of CAZy families were coherent to pathways of degradation of alginate, FCSP, laminarin, carrageenan, starch and agarose (Table S8). Further, CAZymes present within classified CGCs were used as templates to identify non-CGC associated CAZymes that contained similar features (e.g., CAZy family, EC similarity, Pfam and TIGRfam annotations). Features frequently occurring within these CGCs (top third most frequent annotation), were used to filter genes associated with alginate, laminarin, FCSP and carrageenan pathways (Table S10).

**SUPPLEMENTARY FIGURES**

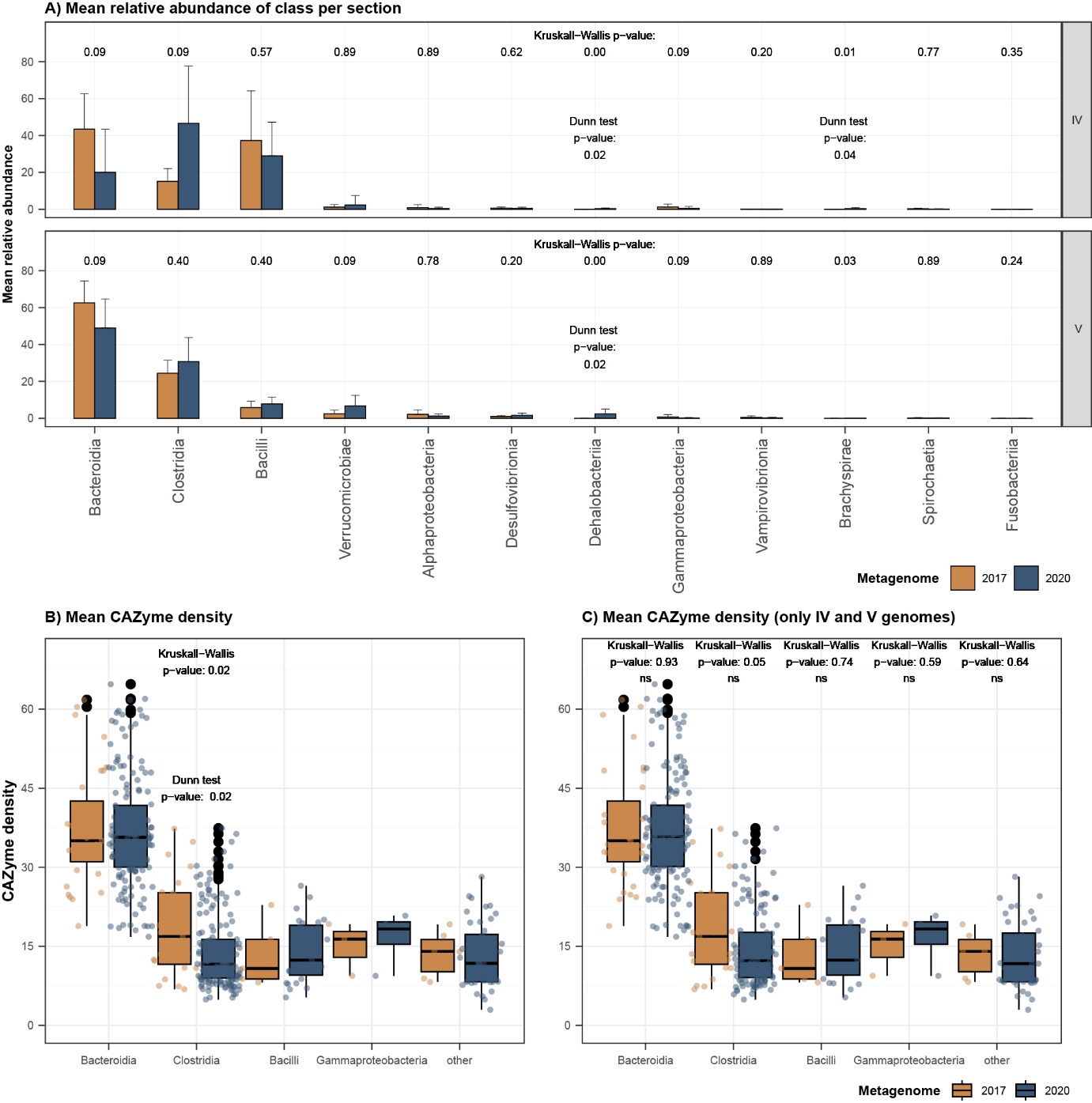

**Figure S1 Mean relative abundance and CAZyme density comparison with a larger metagenome dataset (10 individuals).** (A) Bar plots of mean relative abundance per gut section and bacterial class across metagenomes assembled from fish collected in 2017 (four fish, 68 unique MAGs; yellow bars) and 2020 (six additional fish collected in summer 2020, 397 unique MAGs available under same BioProject; blue bars). (B, C) Boxplots displaying the CAZyme density of Bacteroidia, Clostridia, Bacilli, and Gammaproteobacteria classes from the 2017 and 2020 datasets. Classes containing less than three MAGs in the metagenome of 2017 are displayed as “other”. The boxes and central line represent the interquartile range and median CAZyme density across MAG in the class, and the whiskers represent the minimum and maximum values within 1.5 times the interquartile range. (C) Boxplots displaying CAZyme density per class omitting MAGs recovered from section III in 2020.

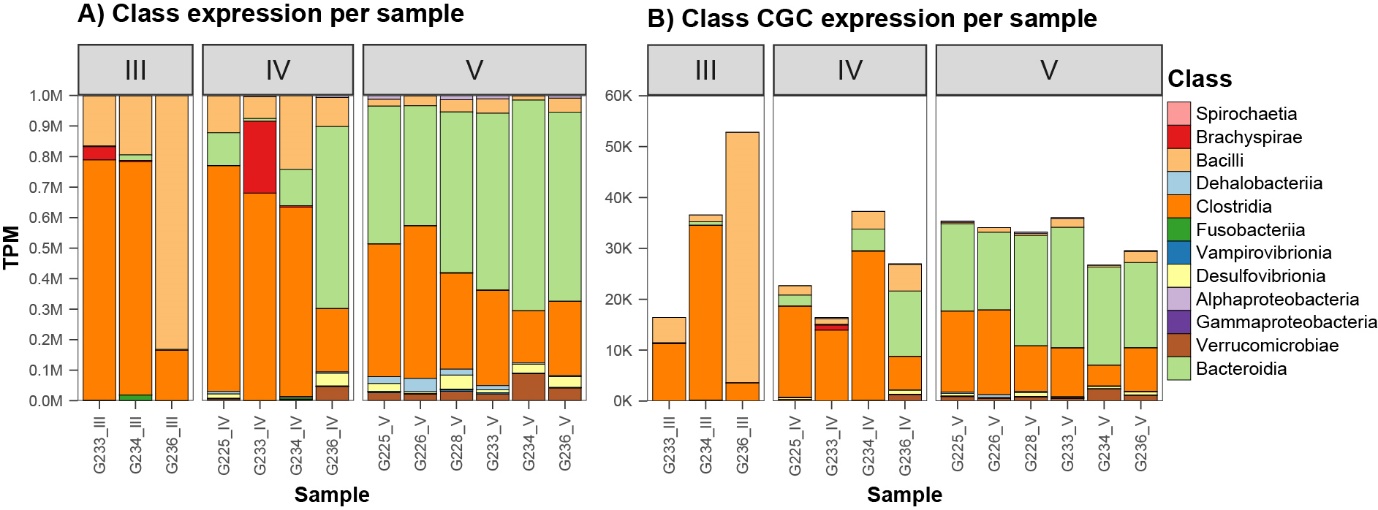

**Figure S2 Gene expression of bacterial classes across sample.** (A) Bar plots showing overall gene expression by class per sample in gut sections III, IV and V. Bars are coloured according to class. Y-axis show the summed expression per sample of each class (labels on the left) and x-axis indicate the sample (fish identifier and gut section). TPM = transcripts per million. (B) Stacked bar plots showing the summed expression of CGC genes per class across samples.

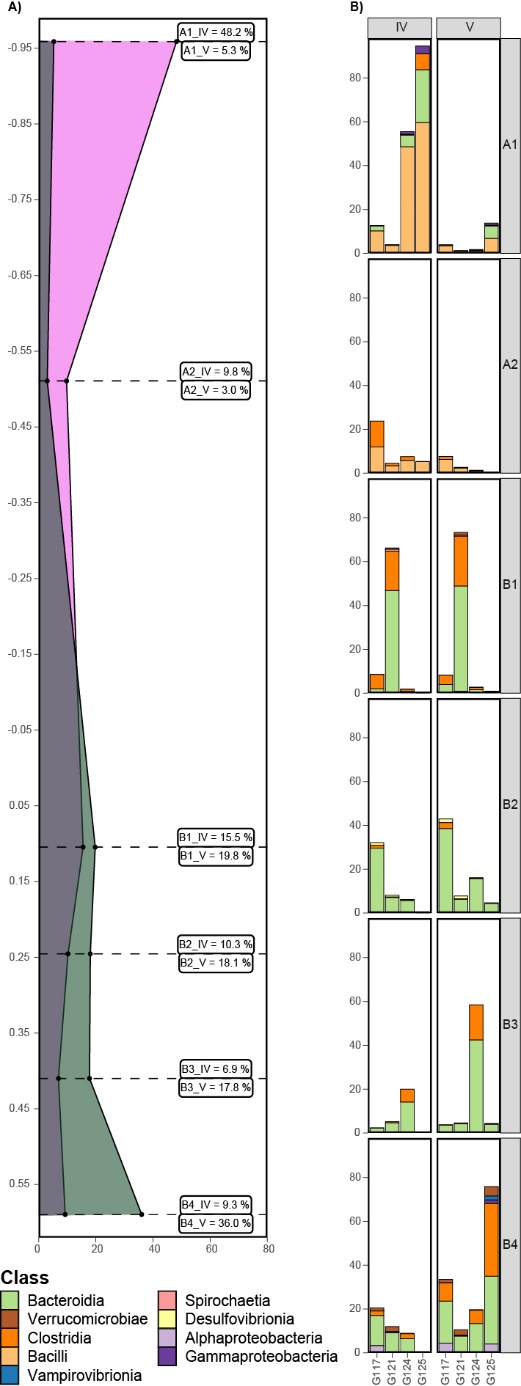

**Figure S3 Longitudinal distributions of co-abundant taxa.** (a) Line plot displaying the summed relative abundance of each co-abundant group in sections IV and V (x-axis and inner labels), and their Log10 fold changes from section IV to V (y-axis). Shading denotes section relative abundance in sections IV (pink) and V (dark green). (b) Stacked bar plots show the relative abundance (%) of each co-abundant group (A1-B4) across individual fish (G117, G121, G124, G125) and gut section (IV, V). Bar colours denote class.
